# Wildlife feeding increases risk of male wild turkeys (*Meleagris gallopavo*) to hunter harvest

**DOI:** 10.64898/2026.04.30.721985

**Authors:** Marcus Lashley, Elizabeth Leipold, Brandon McDonald, Carolina Baruzzi

## Abstract

Wildlife feeding during the wild turkey (*Meleagris gallopavo*) hunting season is legal in many states within the United States, but hunting turkeys with the aid of bait is unlawful in most states. The most common policy to prevent wildlife feeding from acting as bait is to restrict hunting within a defined radius. However, the effect of wildlife feeders on turkey harvest risk and the effectiveness of distance restrictions on mitigating that influence have not been investigated. During 2024–2025, we used GPS transmitters to track 30 adult male turkeys during the spring hunting season on private land with active feeders in Florida, USA, where hunting turkeys within a 91 m radius of a feeder was unlawful. We used Cox proportional hazard models to link risk of hunter harvest with unique feeders visited daily, number of feeders within a home range, and average morning distance and roosting distance to feeders at multiple temporal scales. Hunters harvested 53% of the tagged turkeys. Risk of hunter harvest increased with the number of unique feeders visited the previous day and after the first three days of hunting season with the number of active feeders within a home range. As distance from the most recent roost site and average morning distance to a feeder decreased, risk of hunter harvest increased. We estimated that risk of hunter harvest would be reduced by over 50% if distance restrictions were increased from 100 m to 200 m, by nearly 75% with an increase from 100 m to 300 m, and by nearly 90% with an increase from 100 m to 500 m. To completely eliminate the influence of wildlife feeders on risk of hunter harvest would require a restriction distance well beyond a 500m radius, which is impractical given that this radius would result in an area twice the average private landowner property size in the region. Thus, if wildlife feeding during the turkey hunting season is to be allowed, it will act as bait, in which case, the acceptable level of its influence as bait can be achieved with the appropriate hunting radius restriction.

---

Provisioning food for wildlife is widespread, often through the use of wildlife feeders (Oro et al. 2013, Murray et al. 2016, Bordjan et al. 2023). Wildlife feeders can be defined as stations where food is continually replenished and made available to wildlife from a single point source. They are commonly used for the supplemental feeding and baiting of game species in the United States (U.S.), where feeding or baiting (for at least one species) is legal in 49 of 50 states (Holm 2022). While legal across the U.S., this practice is particularly prevalent across private lands in the southeastern United States (Holm 2022). While the tracking of this practice varies from state to state, recent reports indicate that > 80% of hunters and private landowners in Arkansas, 94% in South Carolina, and 92% in North Carolina hunt over bait or otherwise use wildlife feeders on private lands (Zellers 2019, South Carolina Department of Natural Resources 2020, Jewell et al. 2024). In South Carolina, the Department of Natural Resources estimated that 18 kg/ha/year was placed out by hunting clubs in 2006 (SCDNR 2020), while 206 kg of bait or supplemental feed per licensed hunter per year was provided on the landscape in 2023 in North Carolina (Jewell et al. 2024).

Despite wildlife feeders often being used for a particular species, non-target species frequently take advantage of these resources (Guthery et al. 2004). The wild turkey (*Meleagris gallopavo*), a prominent game species is one of the most common non-target species found at wildlife feeders (Hensen et al. 2012, Huang et al. 2022a). This raises potential concerns because the effects of non-target feeding on wild turkeys remain largely unknown, and with evidence of population declines in part of its range, there is a need to better understand the role of feeding on wild turkey conservation (Eriksen et al. 2015, Chamberlain et al. 2022). In particular, wildlife feeding can increase the predictability of an individual’s location and movement behavior (Rollins et al 2009, Griffin 2017). Corn (*Zea mays*) has been successfully used as bait when capturing wild turkeys for conservation, translocation, and research suggesting a similar effect on turkey movement (Bailey et al. 1980, Cobb et al. 1995, Delahunt et al. 2010). Moreover, being more predictable may be associated with increased probability of a male turkey being harvested (Gulotta et al. 2025). However, hunting turkeys with the aid of bait is unlawful throughout most of their range and distance restrictions are often imposed in an effort to mitigate the influence of wildlife feeding on risk of hunter harvest during the spring turkey season such that the wildlife feeding does not act as bait.

Declines in wild turkey populations have prompted many state agencies to evaluate factors that influence male or “gobbler” harvest rates and the effects of spring hunting on populations as a whole (Isabelle et al. 2018, Quehl et al. 2024). While gobbler harvest rates across the Southeast remain relatively stable in landscapes mostly without wildlife feeders (∼30% of adult males; Wightman et al. 2024), it is unknown how risk of individuals to hunter harvest or population level harvest rate is affected by wildlife feeding. Since active wildlife feeders have been demonstrated to concentrate female wild turkey movements, roosting, and affect social behavior during the spring breeding season (Huang et al. 2026), there is strong potential that wildlife feeders could influence the vulnerability of males to hunter harvest during the hunting season even though hunting is usually restricted near the feeders. We hypothesized that wildlife feeders would serve as bait by increasing the risk of turkeys to hunter harvest. Our objectives were to quantify the influence of wildlife feeders on the risk of male turkeys to hunter harvest across various spatiotemporal scales and explore how distance restrictions might affect that influence given that most state agencies have adopted distance restrictions for policy and enforcement.

## STUDY AREA

Our study occurred during the 36-day Florida spring turkey hunting season in 2024 (16 March–21 April) and 2025 (15 March–20 April) on two privately owned properties in south-central Florida, USA (Figure 1), where spring turkey hunting and active use of wildlife feeders by wild turkeys occurred simultaneously. DeLuca Preserve (hereafter DeLuca), was a 11,000 ha ranch located in Osceola County owned and managed by the University of Florida. The other property was privately owned with a similar sized area used by tagged turkeys and was located in Osceola and Brevard Counties (hereafter Brevard). Approximately 30 km apart, both properties were located in the Florida Wildlife Corridor, which is a critical state-level coordinated wildlife conservation initiative that is particularly important to the Osceola subspecies of wild turkey (*M. g. osceola*; Sibiya et al. 2026). Both properties were primarily used for cattle grazing and consisted of a mosaic of dry and wet prairie, pinelands, hardwood hammocks, hardwood swamps, cypress swamps, improved pasture, and citrus groves. Habitat management on both properties included prescribed burning in forested areas and prairies, mowing, and occasional timber harvest.

**Figure 1.**
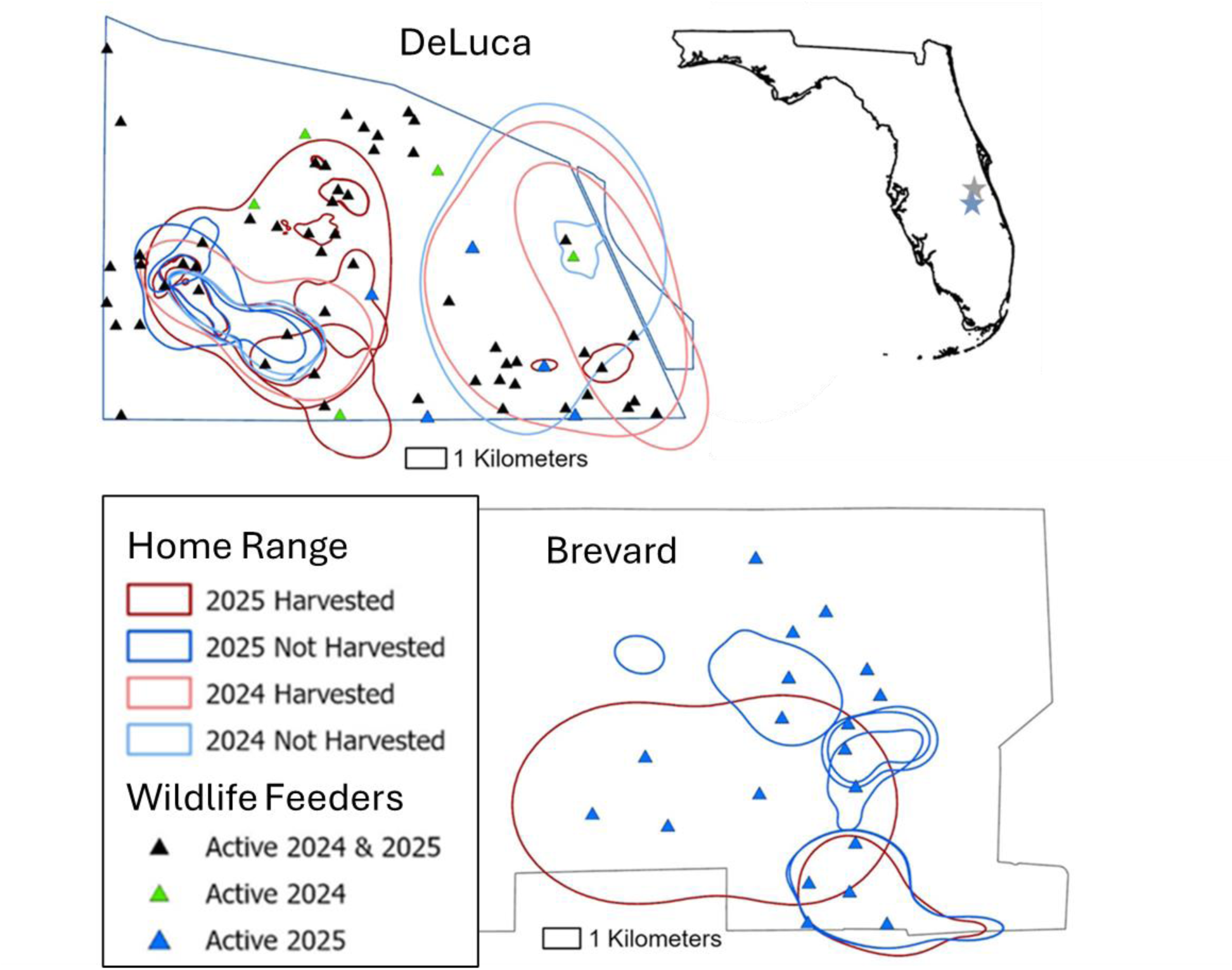
Study area. DeLuca (outlined in blue) and Brevard (outlined in gray). Wildlife feeder locations (triangles) and 95% ranges for harvested and non-harvested turkeys are shown (see text for details regarding calculating 95% range).

Wildlife feeders were active at both properties, replenished with corn year-round. Most feeders were directional spin cast that released food twice per day or gravity released providing access to food *ad libitum*. Field technicians on an ad-hoc basis verified that feeders were active and consistently stocked throughout each hunting season. Feeder locations were recorded using the OnX Hunt application (OnX Maps, Missoula, Montana, United States) on DeLuca and provided by the hunting club on Brevard. In total, DeLuca had 60 active wildlife feeders over both years, though not all feeders in 2024 were active in 2025 and vice versa (Figure 1). Brevard had 19 active wildlife feeders (Figure 1). Distance between a wildlife feeder and the next nearest feeder ranged 140–1,838 m with a mean distance of 589 m on DeLuca and 684–2,182 m with a mean distance of 1,240 m on Brevard.

## METHODS

### Capture and monitoring

We captured wild turkeys between December and February in 2023–2025 using rocket nets in areas temporarily baited with cracked corn. After capture we recorded intrinsic characteristics for each turkey including sex, age (adult or juvenile) as identified by molting pattern in tail rectrices (Williams and Austin 1988), left and right spur length (mm) which we averaged to get a mean spur length, capture mass (kg), and beard length (mm). We banded turkeys with 5/8’’ aluminum rivet bands (National Band & Tag Company, Newport, KY, USA) and attached backpack style GPS (global positioning system) transmitters (custom TechnoSmArt AxyTrex GPS enabled accelerometer with a drop-off mechanism, TechnoSmArt, Rome, Italy, or E-obs Bird Battery 1A internal battery tag models, E-obs, Grünwald, Germany) that stored turkey locations on-board and also provided a VHF (very high frequency) radio signal to assist with tracking. The TechnoSmArt transmitters were programmed to begin recording locations on March 1st until turkey death or on-board memory was full, which generally occurred > 3 weeks after the hunting season had concluded. The TechnoSmArt transmitters recorded hourly locations from 6:00–20:00 EST with roost locations recorded at midnight (0:00 EST) and 4:00 EST in 2024 and at 21:00 EST and 3:00 EST in 2025. Data from the TechnoSmArt transmitters was only available after successful recovery of a transmitter. The e-obs transmitters (deployed only in 2025) began recording locations directly after capture release and data was available immediately via remote download. Data was removed from a transmitters’ on-board data storage after a successful transmission, which allowed data to be recorded for a much longer duration after deployment.

But, once on-board storage capacity was reached, no new locations would be recorded until data was successfully downloaded. We downloaded data 1–2 times per week provided a turkey could be located. The e-obs transmitters recorded locations every 30 minutes from 4:30–21:00 EST and at midnight (0:00 EST) at DeLuca and from 5:00/5:30–20:00 EST at Brevard. Midnight roost locations were not recorded at Brevard until the last week of March. We tracked male turkeys and analyzed movement patterns from 1 March until the end of the hunting season. When turkeys were harvested, date of harvest was provided by the hunter when returning the transmitter. Non-harvest mortality was estimated and dated using location data and verified within 5 days in the field. All turkey-related procedures (i.e. capture, handling, tracking) were approved by the University of Florida Institutional Animal Care and Use Committee (IACUC 202300000010) and permitted by Florida Fish and Wildlife Conservation Commission (permit 11282310).

### Location data

We removed potentially inaccurate (HDOP > 5 for the TechnoSmArt transmitters) and erroneous locations (from visual inspection). We limited our dataset to those turkeys that were alive at the start of the hunting season (16 March 2024 and 15 March 2025), and only included locations between 1 March and end of the hunting season (21 April 2024 and 20 April 2025) or before 6:00 EST on day of harvest. This ensured that locations post-harvest were not included in the analysis as exact times of harvests were unknown. We subsampled locations from the e-obs transmitters to be within 10 minutes of the hour so that the timing of locations were approximately the same between the transmitter types. We classified locations based on time of day as either roosting locations (19:00–6:00 EST) or day locations (6:00–19:00 EST), and then day locations were subdivided into morning locations (6:00–12:00 EST) and afternoon locations (12:00–19:00 EST) to account for changes in hunting pressure throughout the day (Cartwright and Smith 1988, Gerrits et al. 2020). Most turkeys were harvested at the beginning of the hunting season, and sunrise and sunset during this time occurred at 6:30 and 18:30 EST, suggesting that these were reasonable cut-offs for delineating day and roosting time periods.

We examined turkey interactions with feeders over four different temporal scales (i.e. different lengths of time) for different times during a 24-hour period (19:00 the previous day until 19:00 on the current day; Figure 2). The four different temporal scales were 1) most recent roost location which occurred between the previous day and current day on which the event occurred (harvest or no-harvest), 2) previous days behavior (called previous 1 day in analysis), 3) average behavior over the previous 3 days, and 4) average behavior over the previous 7 days (Figure 2). The four temporal scales allowed us to represent both variation in potential hunter scouting behavior (i.e., only scouting the day before, or multiple scouting events over the previous 3 or 7 days before a hunt), levels of consistency in turkey behavior (e.g., repeated visitation to a location or area), and whether a turkey moving closer or farther to a feeder as they leave the roost on the day of the hunt affected harvest risk. We calculated the average minimum distance to an active feeder for each time of day (roost, day, morning, afternoon) for each temporal scale. For the purposes of the “unique feeders visited” variable we considered a feeder visit to be a location within 200 m (average median distance between hourly locations during the day; Table S1, available in Supporting Information) and used those feeder visits to calculate the total number of unique feeders visited in a day for three of the temporal scales (previous 1, 3, and 7 days).

**Figure 2.**
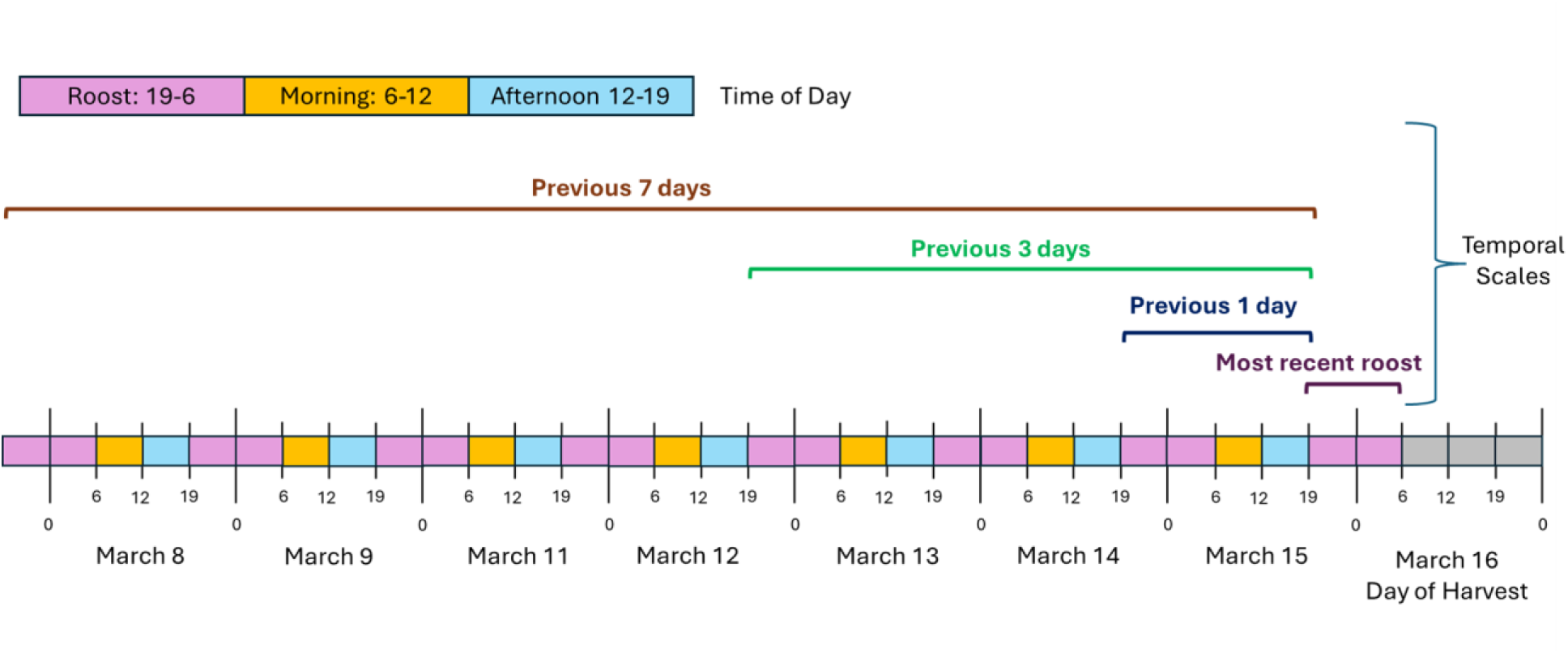
Diagram illustrating the different times of day (roost, morning, and afternoon) and the four temporal scales (most recent roosting, average over the previous 1 day, average over the previous 3 days, and average over the previous 7 days) evaluated within our models.

### Feeders per home range

We calculated the 95% range for each individual using location fixes from 1 March–end of the hunting season or last known alive location (whichever came first) using autocorrelated kernel density estimation using the *ctmm* and *move* packages in program R (Fleming and Calabrese 2025, Kranstauber et al. 2024, R Core Team 2024). We used the “ctmm.select” function to select the best process for modeling the movement data for each individual (Fleming and Calabrese 2025), with Ornstein-Uhlenbeck Foraging (OUF) anisotropic process selected 51% of the time, Ornstein-Uhlenbeck (OU) anisotropic selected 37% of the time, Ornstein-Uhlenbeck (OU) selected 6% of the time, and Ornstein-Uhlenbeck Foraging (OUF) selected 6% of the time. We estimated the percentage of each individuals’ 95% range outside its relative study area and excluded individuals with > 50% of their range outside their relative study area from the analysis as adjacent properties may have unknown feeder locations and different hunting pressure. We then calculated the number of different feeders within each 95% range.

### Harvest risk analysis

We evaluated the effects of wildlife feeders on the risk of hunter harvest for male wild turkeys with Cox proportional hazard models using the “coxph” function in the *survival* package in program R (Therneau 2024, R Core Team 2024). The cox proportional hazard model is a semi-parametric time-to-event model where hazard risk 𝜆_𝑖_(𝑡) (e.g. risk of harvest) is modeled as a function of a non-parametric baseline hazard 𝜆_0_(𝑡) and the parametric covariates *X* (e.g. intrinsic characteristics and different measurements of feeder use) that affect hazard risk, and was specified for all times (𝑡) for turkey 𝑖 as

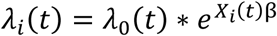

where positive𝛽￼ coefficients were associated with greater risk of harvest and negative coefficients were associated with reduced risk (Therneau and Grambsch 2000). The resultant hazard ratios (HR),𝛽￼), represent the relative change in risk between different values for a covariate where HR > 1 represents increased risk with the change in covariate value, < 1 represents reduced risk, and 1 represents no change in risk (Therneau and Grambsch 2000). Cox proportional hazard models can incorporate both time-independent (fixed) covariates and time-dependent covariates (e.g., minimum distance to a feeder throughout the hunting season) where different time intervals can be constructed for each individual that align with the change in covariate value (Therneau and Grambsch 2000, Therneau et al. 2024). A key assumption of Cox proportional hazard models is that the relationship between a covariate and relative change in hazard is consistent over time, and if this assumption is violated then additional model specifications may be required (e.g. time-varying effects; Therneau and Grambsch 2000, Therenau et al. 2024).

We first conducted a preliminary analysis using time-independent data and univariate models to evaluate the effect of intrinsic characteristics (capture mass, beard length, and average spur length) on harvest risk and then included the most influential intrinsic characteristic (i.e., capture mass, Table S2) in a series of analyses evaluating the effects of feeders on harvest risk using both time-independent and time-dependent variables. We defined an event as either harvested (event = 1) or non-harvested (event = 0), which included those turkeys that survived the hunting season, were predated, or had unknown status due to transmitter malfunction or the turkey migrating outside the study area, etc. For models fitted with multiple covariates, we assessed correlation using Spearman’s correlation coefficient. We considered a correlation of > |0.70| to be highly correlated and found no strong correlation between any variables. We evaluated the proportional hazard assumption using Schoenfeld residuals and formal tests of whether the time-dependent coefficient had a slope of 0 and considered the proportional hazard assumption to be violated if P < 0.05 (Therneau and Grambsch 2000, Therneau 2024, Therneau et al. 2024, R Core Team 2024). If the proportional hazard assumption was violated for a covariate within a model, we incorporated a time-varying coefficient by dividing time into intervals and modeling the effect as a piecewise constant that assumed the hazard ratio was constant within each time interval (Therneau et al. 2024, Therneau 2024, R Core Team 2024). To account for observations being correlated within study sites, we computed a robust variance using the Huber sandwich estimator (Therneau and Grambsch 2000). We evaluated whether the model was considered different from the null (P < 0.05) using the Wald test (Therneau and Grambsch 2000) and model fit with the concordance statistic (i.e., how well the model predicts which event occurred first within a pair of events), where we considered a value > 0.60 to represent acceptable model fit (Therneau and Atkinson 2024). We used Efron estimation to handle ties (Therneau and Grambsch 2000).

We constructed two different datasets: one with only time-independent data, and a second with time-independent and time-dependent data to account for differences in movement or behavior on a day-to-day basis. For the dataset with only time-independent data, time-to-event was defined as the start of the hunting season (day 0) until hunter harvest (event = 1). Individuals that were not harvested were right-censored (event = 0) at the end of the study (day 36) or last known date alive if predated or no longer able to be located. For the second dataset with both time-dependent and time-independent variables, we constructed daily time intervals beginning at the start of the hunting season and ending at harvest. Unharvested individuals were right-censored at either the end of the study (interval day 35 to day 36) or last known date alive with movement data if the transmitter failed before harvest, an individual was predated, or an individual was no longer able to be found. Because a time interval of (0, 0) cannot be used, for individuals who were harvested on the first day of the hunting season, the time interval was recorded as (0, 0.5; Therneau et al. 2024).

We evaluated the effects of feeders on harvest risk in three contexts: 1) number of feeders within a 95% range (time-independent data), 2) total number of unique feeders visited daily (time-dependent data), and 3) minimum distance to a feeder while roosting and during morning foraging (time-dependent data). To evaluate the effect of number of feeders within a 95% range, we fit a single model with capture mass and number of feeders within a 95% range as fixed effects. To evaluate the effect of the number of unique feeders visited daily, we fit three different models, each with capture mass and average number of unique feeders visited daily over one of three different temporal scales (i.e. previous day, average previous 3 days, and average previous 7 days). To evaluate the effect of minimum distance to a feeder on harvest risk we fit four models, each with capture mass and minimum distance over one of four temporal scales (i.e. most recent roosting, previous day, average previous 3 days, and average previous 7 days). For the models evaluating the effect of minimum distance to a feeder, we included minimum distance for two time of day periods: roost (19:00– 6:00), and morning (6:00–12:00). The model evaluating the effect of most recent roost only had the roost time period. We chose these times of day because most turkey hunting occurs during morning hours which encompasses turkeys coming off the roost and morning foraging (Cartwright and Smith 1988, Gerrits et al. 2020).

Within each model, we log-transformed minimum distance for both roost and morning foraging variables to allow for a diminishing relationship between distance and mortality risk, as we assumed the effect of feeders would decrease with increased distance.

To quantify whether the feeders had a positive or negative effect on the hazard risk, we examined the beta coefficients and hazard ratio. If the 95% confidence intervals for the hazard ratio overlapped 1 then we did not consider there to be a statistically significant relationship. To evaluate the effect of different distance restrictions we evaluated the hazard ratio between 100 m (approximately 100 yards, which was the concurrent distance restriction in Florida) from a feeder and increasing distance increments of 100 m (e.g. between 100 and 200 m, 100 and 300 m, etc.). We also evaluated changes in risk in increasing 100 m intervals (e.g. between 100 and 200 m, 200 and 300 m, 300 and 400 m, etc.).

## RESULTS

We deployed a total of 52 GPS transmitters during the study period. In 2024, we deployed 11 GPS transmitters (all TechnoSmArt) on male turkeys at DeLuca and in 2025 we deployed 17 GPS transmitters (all e-obs) at Brevard and 24 GPS transmitters at DeLuca (9 e-obs and 15 TechnoSmArt). We removed a total of 22 turkeys from the analysis: 10 due to loss before the hunting season began (e.g., capture myopathy, dropped transmitters, inability to relocate after capture, or death before the hunting season began), 1 because intrinsic characteristics were not recorded and were therefore automatically removed while fitting the model, 4 because over 50% of their home ranges were located outside their respective study areas, and 7 due to problems obtaining data from the transmitters (e.g., transmitter was damaged before recovery, transmitter was unrecoverable, malfunction of onboard data storage). We had a total of 30 male turkeys in the final analysis (7 at Deluca in 2024, 15 at DeLuca in 2025, and 8 at Brevard in 2025), with a total of 16 harvested (3 at DeLuca in 2024, 11 at DeLuca in 2025, and 2 at Brevard in 2025). We had 6 turkeys harvested on the first day of the hunting season, 2 on the sixth day, and 1 on days 3, 5, 7, 10, 11, 18, 21, and 28 of the hunting season. Of the non-harvested male turkeys, we had 2 with unknown fates, 1 that died through non-human predation, and 11 that were verified alive at the end of the hunting season. During 2025, the transmitter of one male died 10 days before harvest (day 28) causing a loss of a minimal amount of movement data that resulted in the turkey being censored on day 18 during the analyses examining the effect of total number of unique feeders visited daily and minimum distance to feeder (resulting in 15 harvest events versus 16).

### Effects on risk of hunter harvest

#### Intrinsic characteristics

Intrinsic characteristics (e.g., capture mass, beard length, and mean spur length) did not vary between properties (Figure S3). On average capture mass was smaller and beard length was shorter in hunter harvested than non-harvested male turkeys (Figure S3). Of the three univariate models examined (capture mass, beard length, mean tarsus length), only the model with capture mass was different from the null model (Wald test, P < 0.01) and was therefore used in all the subsequent models evaluating the effects of feeder use on harvest (Table S2). In the univariate model and all subsequent models, capture mass had a negative relationship with hazard rate (i.e., lower capture mass was associated with increased risk of hunter harvest; Tables S3, S4).

#### Number of feeders within home range

The number of feeders within a home range varied from 2–22 at DeLuca and 3–5 at Brevard, but the median number of wildlife feeders within a home range was similar (4.0 and 4.5 respectively, Table S5). While home ranges had greater variability in size at DeLuca (1.1–77.9 km^2^) versus Brevard (7.2–28.7 km^2^), median home range at DeLuca (3.6 km^2^) was smaller than Brevard (12.6 km^2^; Table S5). The effect of the number of feeders on the risk of hunter harvest changed throughout the hunting season resulting in 3 time intervals being modeled: 0–3 days, 3–8 days, 8–end of hunting season (Figure 3, Table S6). During the initial 3 days of the hunting season, number of feeders within a home range had a weak negative effect on harvest risk (HR = 0.93, 95% CI = 0.87–0.998, P = 0.046). But, after the initial first three days the direction of relationship changed and there was a strong positive relationship (p < 0.001) between the number of feeders within a home range and hazard risk (days 3–8, HR = 1.25, 95% CI: 1.12–1.40), with the effect becoming greater after the first week in the season (days 8–36, HR = 2.49, 95% CI: 2.13–2.90; Figure 3, Table S6).

**Figure 3.**
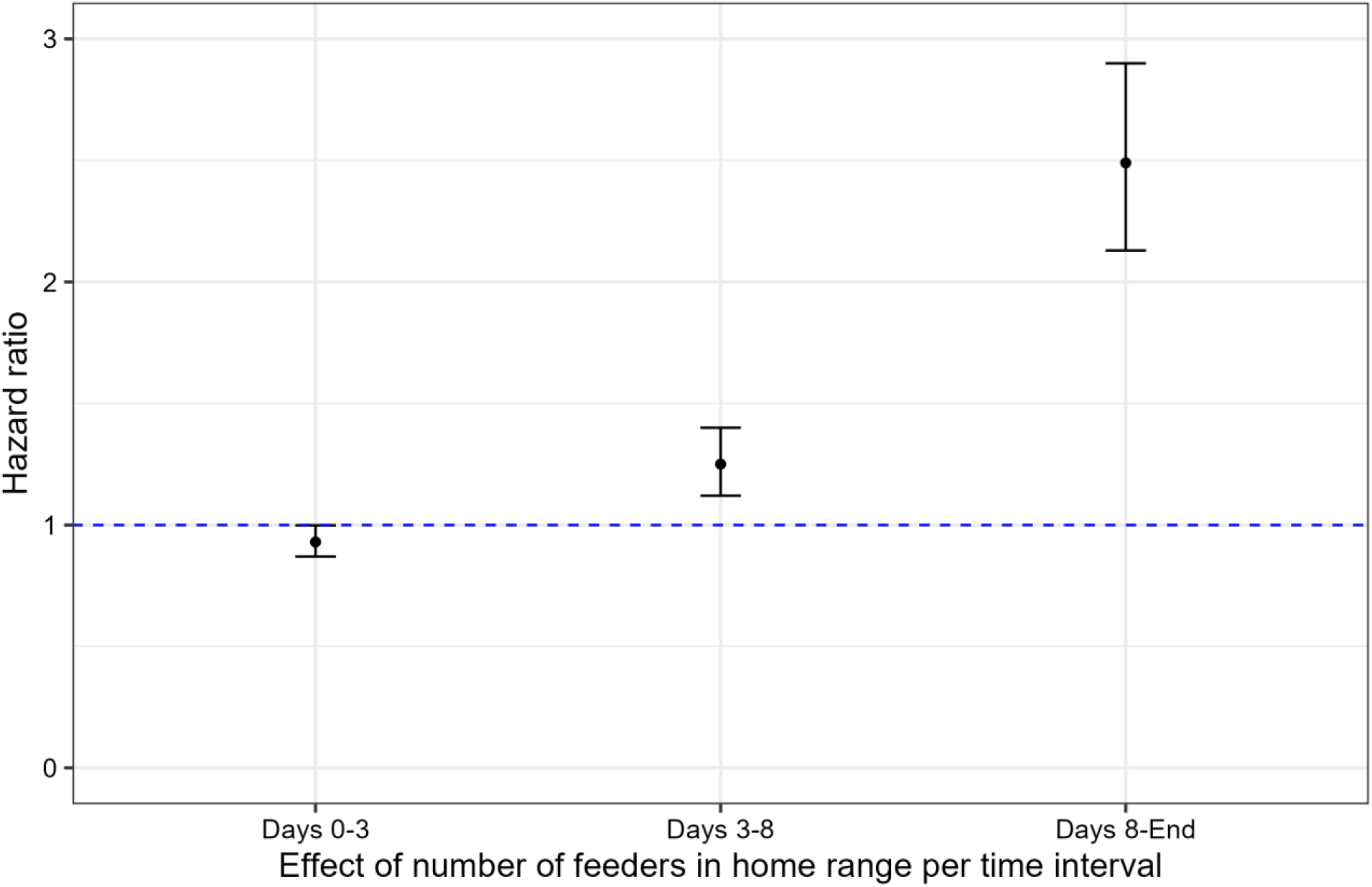
Hazard ratios with 95% confidence intervals for male wild turkey for the effect of number of wildlife feeders within a 95% home range on hazard rate (hunter harvest) in southcentral Florida during the spring hunting season in 2024 and 2025. The model evaluated included capture mass (constant over time), and three different time intervals during the spring hunting season for the effect of number of feeders: days 0–3, days 3–8, and days 8–end of the hunting season. Hazard ratios below 1 (the blue dotted line) indicate a negative relationship with hazard risk while those above 1 indicate a positive relationship with hazard risk (i.e., lower survival).

#### Number of unique feeders visited

The effect of number of unique feeders visited daily was consistent throughout the hunting season (i.e., proportional hazard assumption was met without additional model specifications). On the shorter temporal time scale (previous day), the risk of hunter harvest increased with the number of unique feeders visited daily (HR = 1.85, 95% CI: 1.20–2.85). With longer temporal scales (average wild turkey behavior over the previous 3 and 7 days), unique feeders visited daily and hazard risk were not associated (Figure 4, Table S6).

**Figure 4.**
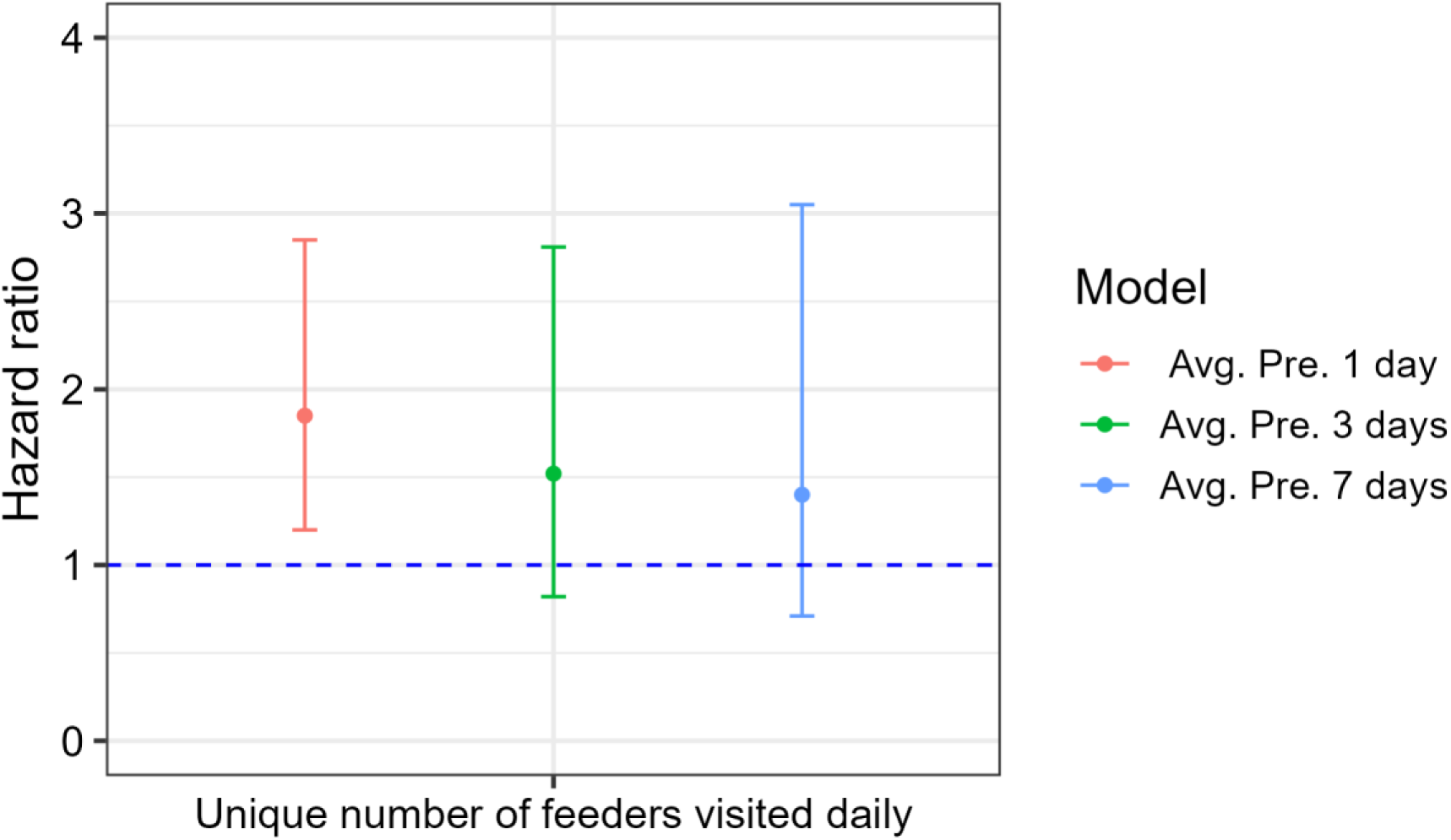
Hazard ratios with 95% confidence intervals the effect of total number of unique feeders visited daily on the hazard rate (hunter harvest) for male wild turkey in southcentral Florida during the spring hunting season in 2024 and 2025. Three models were evaluated: 1) capture mass (constant over time) and average number of unique feeders visited daily over the previous day (varied over time), 2) capture mass and average number of unique feeders visited daily over the previous 3 days, and 3) capture mass and the average number of unique feeders visited daily over the previous 7 days. Hazard ratios below 1 (the blue dotted line) indicate a negative relationship with hazard risk while those above 1 indicate a positive relationship with hazard risk (i.e. lower survival).

#### Minimum distance to a feeder

Minimum distance to a feeder during the morning and most recent roost was strongly related to hazard risk, and the strength of that effect did not change with the progression of the hunting season. During the morning, as average distance to a feeder increased for the previous day (HR = 0.64, 95% CI: 0.62–0.66), previous 3 days (HR = 0.42, 95% CI: 0.41–0.43), and previous 7 days (HR = 0.30, 95% CI: 0.24–0.36), hazard rate decreased (Figure 5, Table S6). While decreasing distance of the most recent roost to a feeder was associated with increased risk of harvest during the morning of a hunt (HR = 0.45, 95% CI: 0.37–0.55), no relationship was observed for the other three temporal scales between distance from a roost to a wildlife feeder and harvest risk suggesting that only the most recent roost site is important in determining the influence of feeder on risk of hunter harvest (Figure 5, Table S6).

**Figure 5.**
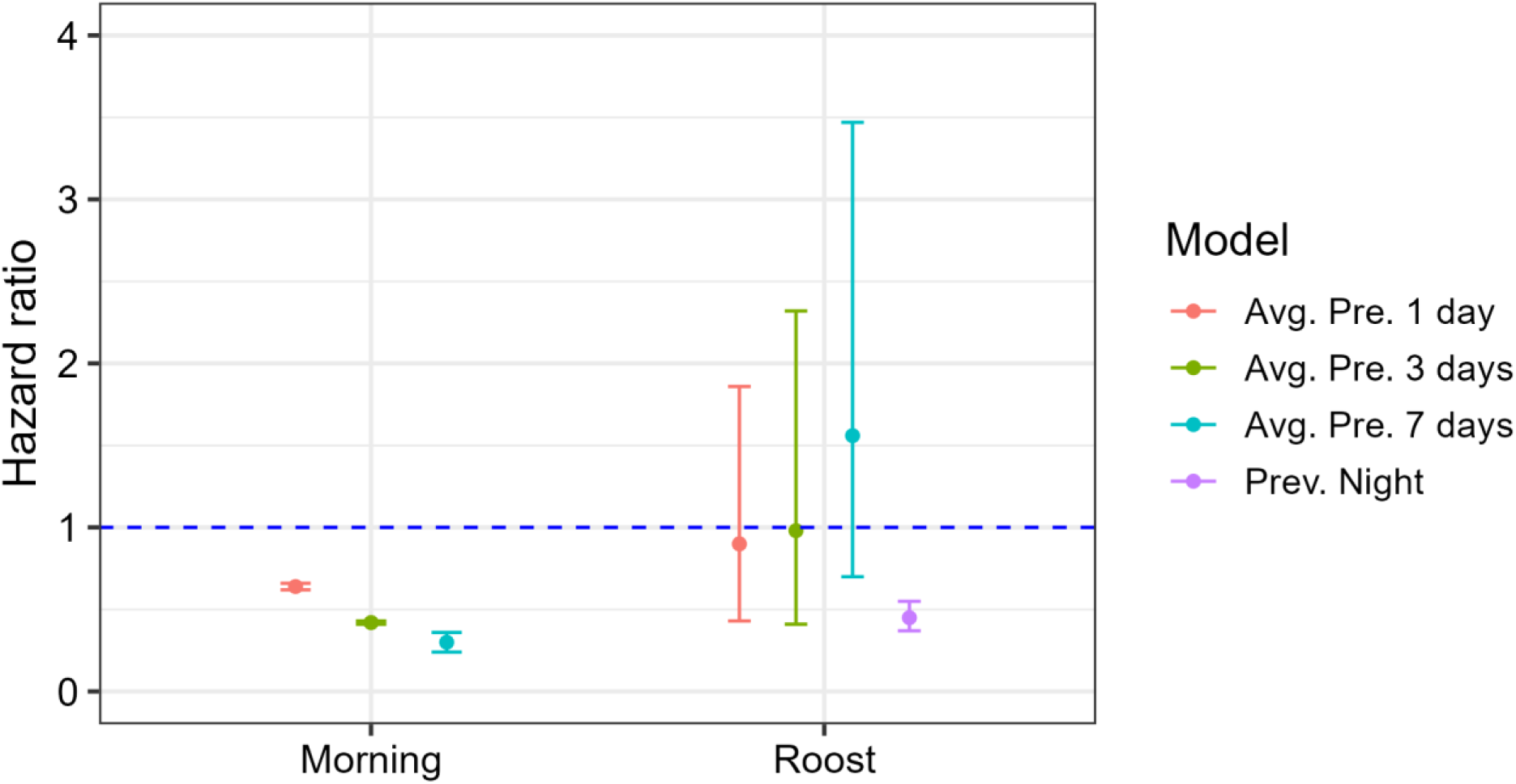
Hazard ratios with 95% confidence intervals for the effect of minimum distance during the morning and while roosting to a wildlife feeder on hazard rate (i.e., hunter harvest) of male wild turkey in southcentral Florida during the spring hunting season in 2024 and 2025. Four models were evaluated: 1) capture mass (time-independent) and minimum distance from a feeder to most recent roosting location (time-dependent), 2) capture mass and minimum distance from a feeder during morning and roosting locations for the previous day, 3) capture mass and average minimum distance from a feeder during the morning and while roosting for the previous 3 days, and 4) capture mass and average minimum distance from a feeder during the morning and while roosting for the previous 7 days. Hazard ratios below 1 (the blue dotted line) indicate a negative relationship with hazard risk for increasing minimum distance to a feeder while those above 1 indicate a positive relationship with hazard risk with increasing minimum distance to a feeder (i.e. worse survival).

When we examined the effect of distance restrictions on harvest risk, changing the distance restriction from 100m to 200m would0.5-fold reduction in hazard risk when considering turkey behavior over the previous week (Figure 6). If distance restrictions were changed from 100 m to 500 m, we observed a > 85% reduction in risk when average turkey behavior over the previous week was considered (Figure 6). The reductions in risk reached a diminishing return beyond 500 m (Figure 6, Figure 7).

**Figure 6.**
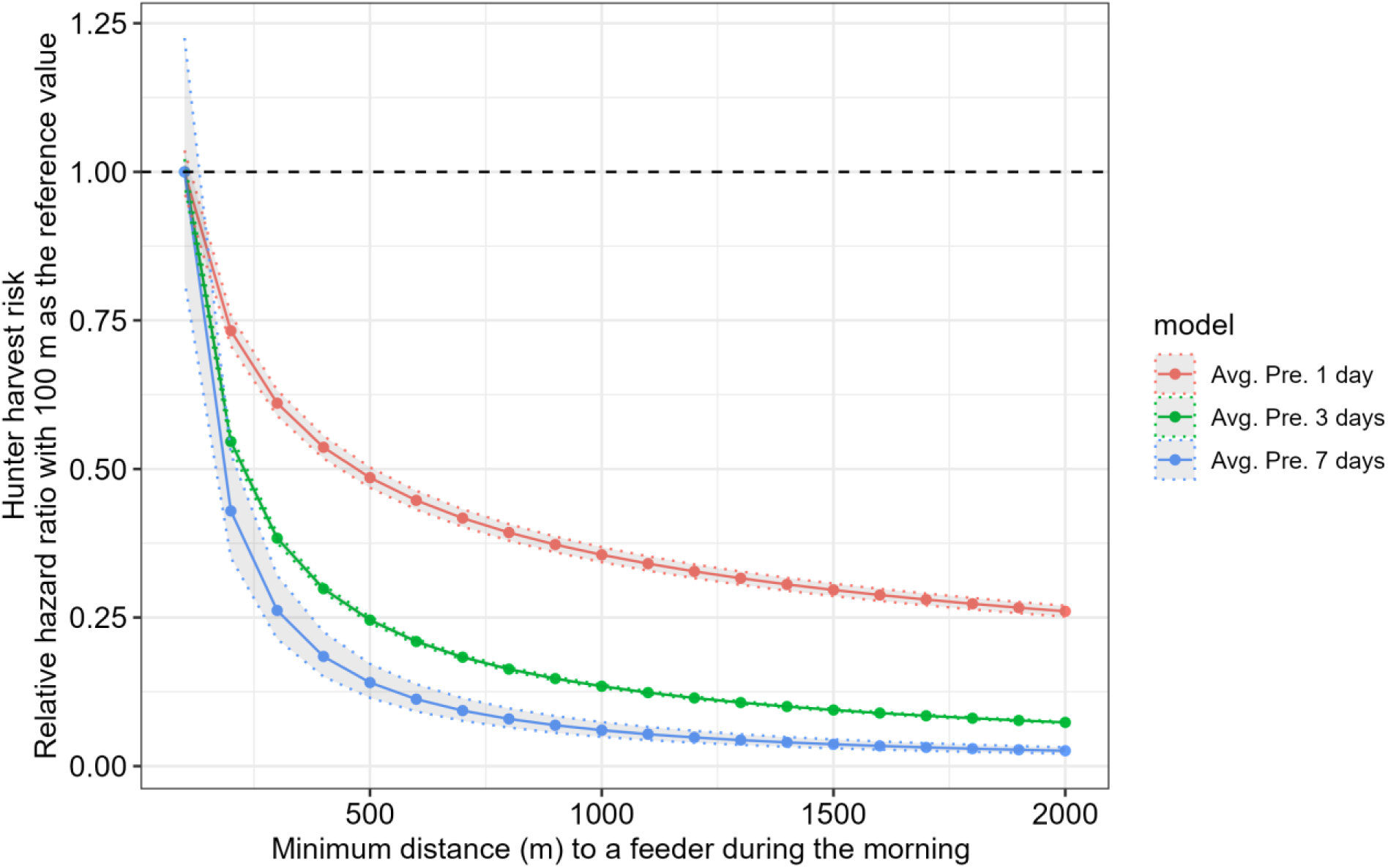
Relative hazard ratios with 95% confidence intervals for an average male wild turkey between 100 m from a feeder (reference) and increasing minimum distance to a feeder during the morning (e.g. between 100 m and 200 m, 100 m and 300 m, 100 m and 400 m, etc.), where the hazard represents the risk of hunter harvest in southcentral Florida during the spring hunting season in 2024 and 2025. Three models were evaluated: 1) capture mass and minimum distance from a feeder during morning and roosting locations for the previous day, 2) capture mass and average minimum distance from a feeder during the morning and while roosting for the previous 3 days, and 3) capture mass and average minimum distance from a feeder during the morning and while roosting for the previous 7 days. Hazard ratios below 1 (the black dashed line) indicate a negative relationship with hazard risk (i.e. higher survival).

**Figure 7.**
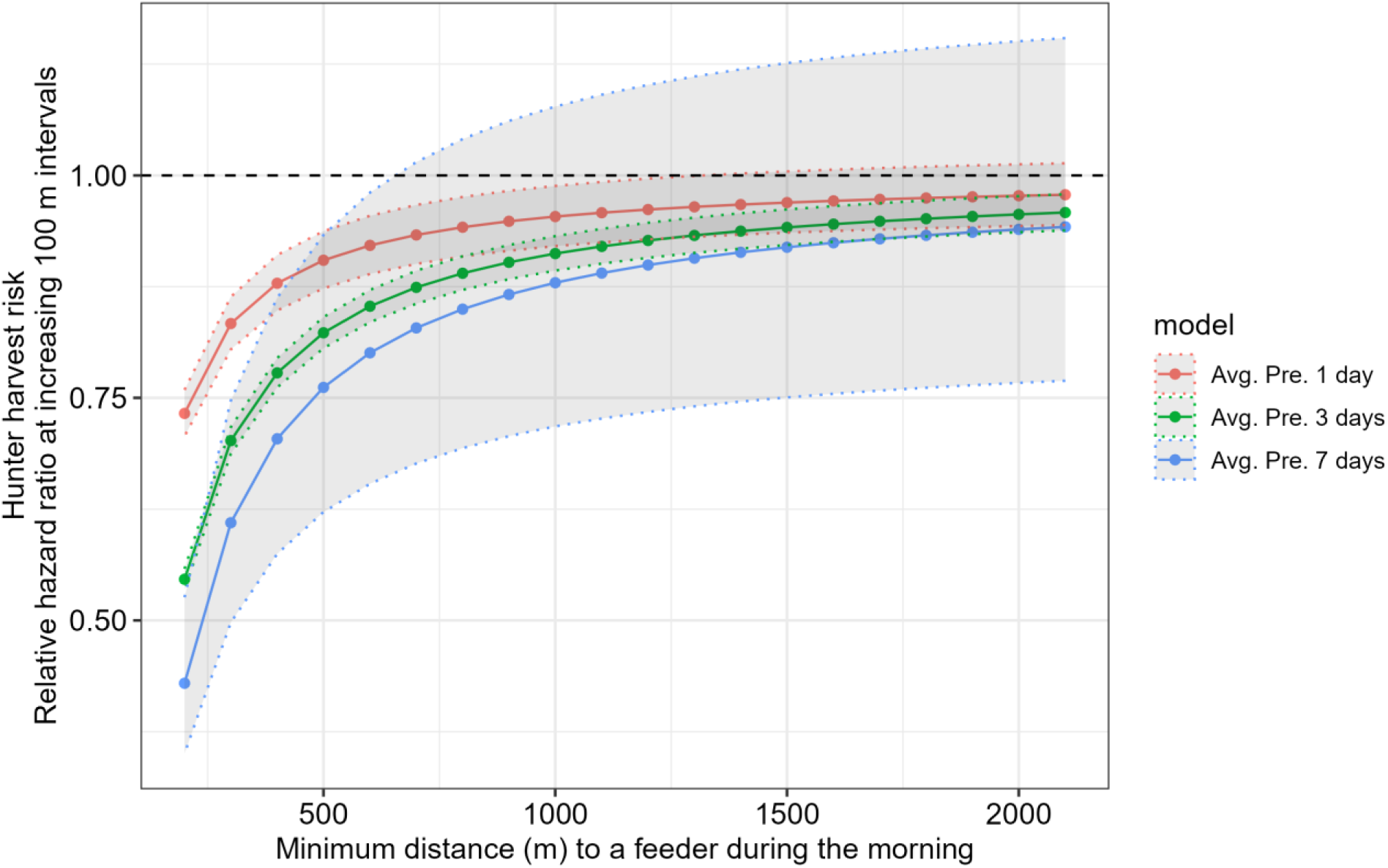
Relative hazard ratios with 95% confidence intervals for an average male wild turkey for increasing 100 m intervals (i.e. first point is the relative hazard ratio between 200m and 100m, the second point is the relative hazard ratio between 300m and 200m, etc.) during the morning, where the hazard represents the risk of hunter harvest in southcentral Florida during the spring hunting season in 2024 and 2025. Three models were evaluated: 1) capture mass and minimum distance from a feeder during morning and roosting locations for the previous day, 2) capture mass and average minimum distance from a feeder during the morning and while roosting for the previous 3 days, and 3) capture mass and average minimum distance from a feeder during the morning and while roosting for the previous 7 days. Hazard ratios below 1 (the black dashed line) indicate a negative relationship with hazard risk (i.e. higher survival). Hazard ratios at 1 indicate that there is no change in hazard between the distances in that interval.

## DISCUSSION

Wildlife feeding increased the risk of hunter harvest for male wild turkeys. The presence of wildlife feeders within a turkeys’ home range, the number of unique feeders visited on a daily basis, the distance between a feeder and roost site, and the distance between a feeder and morning foraging locations were all associated with risk of hunter harvest. These effects were observed across multiple temporal scales for some factors, indicating that both average and most recent behavior affect risk of harvest. Restriction radius, a common enforceable regulation implemented to mitigate the influence of wildlife feeding acting as bait, does affect the degree to which wildlife feeding influences risk of a male turkey to hunter harvest. Our results indicate that a radius far greater than 500 m is required to fully prevent feeders from acting as bait because of a diminishing return of distance that was observed around 500 m.

In the Southeast, most non-industrial private landowner properties were less than 40 ha (Thompson 1996,1997,1999) and property size is decreasing over time (Caputo et al. 2020). If the intent is to not allow bait to influence male turkey harvest, even allowing wildlife feeding on non-hunted properties may also affect legality of hunting on adjacent properties, given the extent of the necessary restrictive radius of > 500 m (an area equivalent to > 78.5 ha) required to eliminate the influence. Some states (e.g., Alabama) have taken the approach to not allow wildlife feeding on the entire property during the turkey season for hunting to be permitted (Alabama Department of Natural Resources 2026). However, even in this case, wildlife feeding on properties that are not actively hunted could still influence the harvest susceptibility of male turkeys on adjacent properties, resulting in a landowner unintentionally participating in unlawful hunting on their own property even though they themselves are not participating in wildlife feeding nor intending to bait. Thus, if wildlife feeding during the turkey hunting season is allowed in any circumstance, it is likely that some level of its influence as bait to turkeys will have to be accepted, and distance restrictions can be established to achieve that acceptable level of influence.

Increased feeder use by individual turkeys probably leads to an increase in risk of harvest because that individual’s roosting and foraging behavior become more predictable. Recently, Gulotta et al. (2025) reported that greater repeatability in male turkey behavior increased the probability of being harvested, and Huang et al. (2026) demonstrated that turkeys that frequented feeders had more predictable movement and roosting behavior.

Given that none of our monitored male turkeys were without access to at least two active feeders, future studies with varying densities of feeders and replication across more study areas would be helpful to fully determine population level effects of feeders, especially on male harvest rate. However, this study represents a real-world scenario of how wildlife feeding is being practiced on private lands in some places. Our observed harvest rate (>50%) was high compared to other studies across the region (∼30%; Wightman et al. 2024). Since feeder proximity and availability influenced susceptibility of male turkeys to hunter harvest, it is plausible that allowing wildlife feeding during the spring turkey hunting season might manifest in a higher male turkey harvest rate, especially since the influence of wildlife feeders is more prevalent later in the season when hunter harvest traditionally has been less concentrated (Wightman et al. 2024). Given that wildlife feeding is one of the most common management practices among private landowners in this region, additional work to understand the ramification for wild turkey conservation and hunting policy is warranted (Jewell et al. 2024).

Quantification of wildlife feeder density is rare. In our study areas, there was 0.5 feeder per km^2^, which is an underestimate of effective feeder density given a large portion of the landscape is not turkey habitat (i.e., open water, developed). It is unknown what the broader effects could be on ecological communities, but chronic nutrient abundance in general simplifies wildlife and plant communities leading to increased abundance of species that are better able to exploit high nutrient availability (Oro et al. 2013). Thus, understanding the magnitude of this practice and the ecological implications is currently an important frontier of ecology. It is likely that chronic availability of this anthropogenic subsidy will have the same effect of simplifying communities over time by benefitting hyper generalist, especially mesomammals that are predators of wild turkeys (Oro et al. 2013). And thus, there is reason for concern not just for wild turkeys but for ecological communities.

Most of the research on wildlife feeding has focused on a number of risks that come with the practices. Wildlife feeding concentrates a suite of species (Lambert and Demarais 2001, Campbell et al. 2013, Candler et al. 2019), which strengthens competition and predation (Oro et al. 2013), increases exposure risk to toxins, and changes intra- and inter-species disease dynamics (Sorensen et al. 2014, Becker et al. 2015, Murray et al. 2016). Wild turkeys commonly use wildlife feeders (Candler et al. 2019) and they are vulnerable to aflatoxins (Quist et al. 2000, Rauber et al. 2007), which often are associated with wildlife feeders at levels known to be toxic to turkeys (Huang et al. 2022a, Oberheu and Dabbert 2001, Schweitzer et al. 2001, Fischer et al. 1995, Dunham et al. 2017, Dale et al. 2025). Also, wildlife feeders are associated with high environmental loading of parasites such as *Eimeria spp.* (Huang et al. 2022b), which can cause coccidiosis in wild turkeys and coccidia infection interferes with mate selection by negatively affecting secondary sexual characteristics in males (reduces iridescence in plumage, Hill et al. 2005, suppresses snood length and skullcap size, Buccholz 1995). Because wild turkeys are ground nesters, they are also potentially vulnerable to hyper-predation during nesting as indicated by artificial nest studies (Cooper and Ginnett 2000; Selva et al. 2014) but could also be vulnerable at other life stages since wildlife feeders subsidize and concentrate relevant predators at those life stages (Oro et al. 2013). Our study shows additional risks of wildlife feeders by illustrating how baiting during the spring turkey season results in increased risk of harvest. Given that wild turkey productivity (Byrne et al. 2015), adult female survival (Lashley et al. 2025), and population size (Casalena et al. 2015, Eriksen et al. 2015, Chamberlain et al. 2022; Londe et al. 2023) declined in the southeast in the mid-2000s concurrent with wildlife feeding becoming common (Jewell et al. 2024), it is plausible that trends in wildlife feeding may have been associated with observed trends in wild turkey populations.

## MANAGEMENT IMPLICATIONS

Our data provides evidence that wildlife feeding serves as bait aiding in the harvest of wild turkeys. The influence of wildlife feeding on harvest risk can be mitigated by radius restrictions on hunting, but eliminating its influence with radius restrictions is not likely a practical option. If wildlife feeding is to be allowed during the spring turkey hunting season it will act as bait, and the level of baiting influence on harvest deemed acceptable must be discussed by state agencies and then can be achieved with the appropriate radius restriction. Given the broad impact of wildlife feeding on ecological communities, future research is also needed to understand how wildlife feeding affects survival during other life stages and seasons (e.g. female adult survival and reproductive vital rates) to fully understand the implications of this practice for the conservation of wild turkeys.

