## Supporting Information for "Wildlife feeding increases risk of male wild turkeys (*Meleagris gallopavo*) to hunter harvest"

S1. Table. Distance between steps during the day (6:00 EST–20:00 EST) from 1 March (approximately 2 weeks before hunting season began)–end of hunting season or mortality. Id = individual, property refers to study site, n is the number of locations used when calculating median and average daytime movement (m). SD = standard deviation.

| ID | Property | Year | Median daytime movement (m) | Average daytime movement (m) | SD daytime movement | n |
|---|---|---|---|---|---|---|
| 1 | DeLuca | 2025 | 187 | 325 | 355 | 167 |
| 2 | DeLuca | 2025 | 146 | 215 | 216 | 369 |
| 3 | DeLuca | 2025 | 254 | 377 | 375 | 583 |
| 4 | DeLuca | 2025 | 100 | 150 | 154 | 203 |
| 5 | DeLuca | 2025 | 150 | 222 | 229 | 384 |
| 6 | DeLuca | 2025 | 299 | 461 | 490 | 237 |
| 7 | DeLuca | 2025 | 135 | 209 | 216 | 296 |
| 8 | Brevard | 2025 | 193 | 317 | 376 | 607 |
| 9 | Brevard | 2025 | 198 | 317 | 378 | 608 |
| 10 | Brevard | 2025 | 234 | 324 | 352 | 423 |
| 11 | Brevard | 2025 | 281 | 371 | 289 | 610 |
| 12 | Brevard | 2025 | 271 | 338 | 273 | 417 |
| 13 | Brevard | 2025 | 204 | 282 | 275 | 287 |
| 14 | Brevard | 2025 | 377 | 499 | 413 | 431 |
| 15 | Brevard | 2025 | 257 | 322 | 300 | 605 |
| 16 | DeLuca | 2024 | 198 | 261 | 248 | 507 |

| | | | | | | |
|---|---|---|---|---|---|---|
| 17 | DeLuca | 2024 | 209 | 307 | 338 | 234 |
| 18 | DeLuca | 2024 | 182 | 301 | 352 | 153 |
| 19 | DeLuca | 2024 | 221 | 323 | 333 | 229 |
| 20 | DeLuca | 2024 | 259 | 369 | 378 | 527 |
| 21 | DeLuca | 2024 | 150 | 231 | 262 | 152 |
| 22 | DeLuca | 2025 | 205 | 279 | 243 | 139 |
| 23 | DeLuca | 2025 | 142 | 213 | 236 | 515 |
| 24 | DeLuca | 2025 | 221 | 325 | 296 | 477 |
| 25 | DeLuca | 2025 | 252 | 388 | 381 | 183 |
| 26 | DeLuca | 2025 | 148 | 196 | 185 | 136 |
| 27 | DeLuca | 2025 | 176 | 246 | 262 | 472 |
| 28 | DeLuca | 2025 | 111 | 199 | 290 | 172 |
| 29 | DeLuca | 2025 | 245 | 396 | 375 | 130 |
| 30 | DeLuca | 2024 | 270 | 365 | 381 | 505 |

S2. Table  Beta coefficients with robust standard errors (SE) and hazard ratios (HR) with 95% confidence intervals (CI) from three univariate cox-proportional hazard models, each evaluating the effect of an intrinsic characteristic (i.e., capture mass (kg), beard length (mm), average spur length (mm)) of male turkeys on risk of hunter-harvest during the spring hunting seasons in south-central Florida in 2024 and 2025. Property was included as a clustering term within all models to account for non-independence among observations within a site. Proportional hazard (PH) assumptions were evaluated using Schoenfeld residuals. To meet the proportional hazard assumption, beard length was modelled as a time-varying effect with two intervals: < 6 days and ≥ 6 days after the start of the hunting season. Also included are the overall model concordance statistic with standard error (Model concordance (SE)) and results of the Wald test (model p-value) evaluating whether a model differs significantly from the null.

| Model | Covariate | Beta Coef. | Robust SE | p-value | HR (95% CI) | PH assumption met | Model Concordance (SE) | Model p-value |
|---|---|---|---|---|---|---|---|---|
| Capture Mass | Capture Mass | -0.73 | 0.13 | < 0.001 | 0.48 (0.37-0.63) | Yes | 0.69 (SE = 0.02) | < 0.001 |
| Beard Length | Beard Length: 0–6 days | 0.006 | 0.01 | 0.38 | 1.01 (0.99-1.02) | Yes | 0.64 (SE = 0.08) | 0.70 |
| | Beard Length: 6–36 days | -0.05 | 0.01 | < 0.001 | 0.95 (0.94-0.97) | Yes | | |
| Mean Spur Length | Mean Spur Length | -0.01 | 0.03 | 0.74 | 0.99 (0.93-1.06) | Yes | 0.52 (SE = 0.04) | 0.70 |

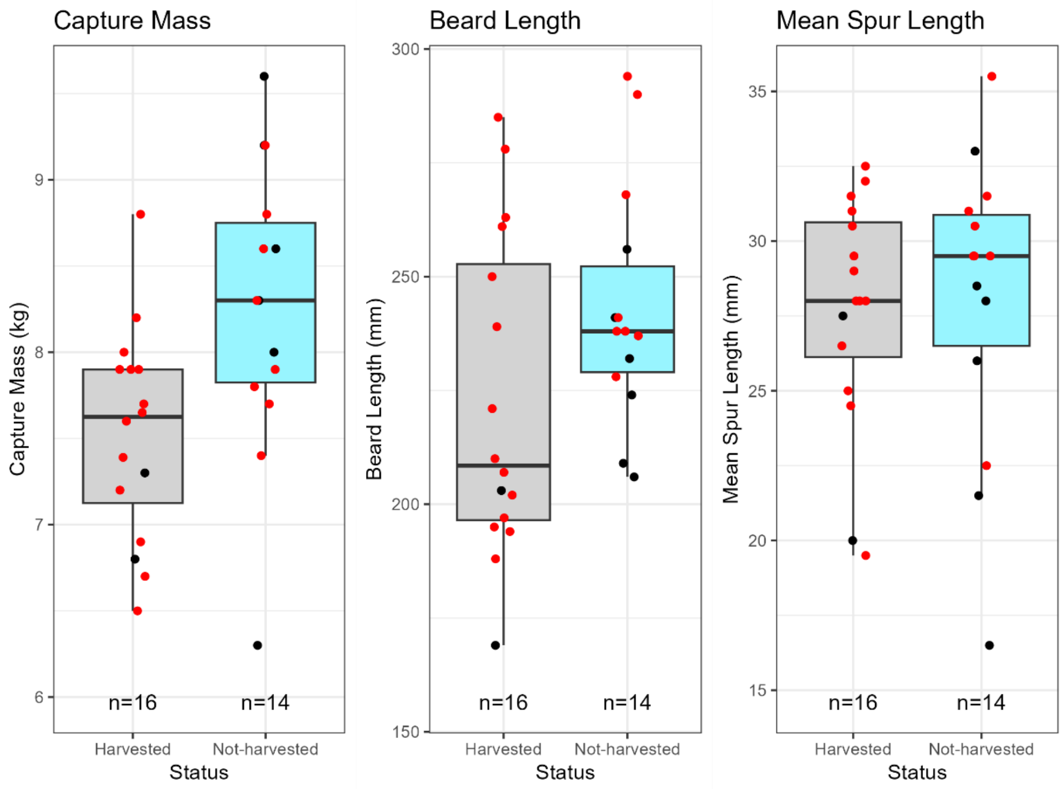


S3. Figure. Box plots of intrinsic characteristics for the harvested (gray) and not-harvested (blue) male turkeys. Points in red represent monitored turkeys at DeLuca and black points represent monitored turkeys at Brevard.

S4. Table. Beta coefficients with robust standard errors (SE) and hazard ratios (HR) with 95% confidence intervals (CI) for capture mass (kg) from a series of cox-proportional hazard models evaluating the effects of capture mass and turkey movement in relation to wildlife feeders on the risk of hunter-harvest for male wild turkeys. Study occurred during the spring hunting seasons in south-central Florida in 2024 and 2025. Models included were the effect of number of feeders within a 95% range, number of unique feeders visited on the previous 1, 3, and 7 days, and the average minimum distance during roosting and morning for the most recent roost and previous 1, 3, and 7 days. Property was included as a clustering term within all models to account for non-independence among observations within a site. MD = minimum distance, Pre 1 = previous 1 day average, Pre 3 = previous 3 day average, Pre 7 = previous 7 day average.

| Model | Covariate | Beta Coef. | Robust SE | p-value | HR 95% CI |
|---|---|---|---|---|---|
| Capture Mass + Number of Feeders in 95% Range | Capture Mass | -0.86 | 0.08 | < 0.001 | 0.42 (0.36-0.50) |
| Capture Mass + Pre 1 Number of Feeders Visited Daily | Capture Mass | -0.9 | 0.04 | < 0.001 | 0.41 (0.38-0.44) |
| Capture Mass + Pre 3 Number of Feeders Visited Daily | Capture Mass | -0.76 | 0.04 | < 0.001 | 0.47 (0.43-0.51) |
| Capture Mass + Pre 7 Number of Feeders Visited Daily | Capture Mass | -0.76 | 0.03 | < 0.001 | 0.47 (0.44-0.50) |
| Capture Mass + Most Recent Roost MD | Capture Mass | -0.91 | 0.002 | < 0.001 | 0.404 (0.403-0.406) |
| Capture Mass + Pre 1 Roost MD + Pre 1 Morning MD | Capture Mass | -0.79 | 0.06 | < 0.001 | 0.45 (0.40-0.51) |

| | | | | | |
|---|---|---|---|---|---|
| Capture Mass + Pre 3 Roost MD + Pre 3 Morning MD | Capture Mass | -0.85 | 0.08 | < 0.001 | 0.43 (0.36-0.51) |
| Capture Mass + Pre 7 Roost MD + Pre 7 Morning MD | Capture Mass | -0.8 | 0.05 | < 0.001 | 0.45 (0.41-0.50) |

S5. The size and number of feeders within a 95% range per individual male turkey. Id = individual and property refers to study site. 95% ranges were calculated using aKDEs.

| Individual | Property | Year | Number of Feeders Within 95% Range | Size of 95% range ($km^2$) |
|---|---|---|---|---|
| 1 | DeLuca | 2025 | 3 | 7.04 |
| 2 | DeLuca | 2025 | 6 | 2.60 |
| 3 | DeLuca | 2025 | 3 | 3.81 |
| 4 | DeLuca | 2025 | 5 | 1.41 |
| 5 | DeLuca | 2025 | 7 | 2.76 |
| 6 | DeLuca | 2025 | 3 | 5.21 |
| 7 | DeLuca | 2025 | 4 | 1.14 |
| 8 | Brevard | 2025 | 4 | 14.93 |
| 9 | Brevard | 2025 | 4 | 14.88 |
| 10 | Brevard | 2025 | 4 | 7.21 |
| 11 | Brevard | 2025 | 5 | 10.38 |
| 12 | Brevard | 2025 | 4 | 28.72 |
| 13 | Brevard | 2025 | 5 | 21.69 |
| 14 | Brevard | 2025 | 3 | 10.40 |
| 15 | Brevard | 2025 | 4 | 7.75 |
| 16 | DeLuca | 2024 | 2 | 3.29 |
| 17 | DeLuca | 2024 | 14 | 33.97 |
| 18 | DeLuca | 2024 | 8 | 13.03 |
| 19 | DeLuca | 2024 | 12 | 30.08 |

| | | | | |
|---|---|---|---|---|
| 20 | DeLuca | 2024 | 3 | 3.48 |
| 21 | DeLuca | 2024 | 8 | 18.59 |
| 22 | DeLuca | 2025 | 3 | 1.21 |
| 23 | DeLuca | 2025 | 5 | 1.55 |
| 24 | DeLuca | 2025 | 5 | 3.39 |
| 25 | DeLuca | 2025 | 22 | 27.10 |
| 26 | DeLuca | 2025 | 4 | 62.71 |
| 27 | DeLuca | 2025 | 5 | 3.64 |
| 28 | DeLuca | 2025 | 2 | 77.94 |
| 29 | DeLuca | 2025 | 2 | 3.55 |
| 30 | DeLuca | 2024 | 3 | 3.53 |

Table S6. Beta coefficients with robust standard errors (SE) and hazard ratios (HR) with 95% confidence intervals (CI) for covariates relating turkey movements to wildlife feeders from a series of cox-proportional hazard models evaluating the effects of capture mass and turkey movement in relation to wildlife feeders on the risk of hunter-harvest for male wild turkeys. Study occurred during the spring hunting seasons in south-central Florida in 2024 and 2025. Models included were the effect of number of feeders within a 95% range, number of unique feeders visited on the previous 1, 3, and 7 days, and the average minimum distance during roosting and morning for the most recent roost and previous 1, 3, and 7 days. Property was included as a clustering term within all models to account for non-independence among observations within a site. Proportional hazard (PH) assumptions were evaluated using Schoenfeld residuals. To meet the proportional hazard assumption, the effect of number of feeders within a 95% turkey range was modelled as a time-varying effect with three intervals: < 3 days, 3–8 days, and ≥ 8 days after the start of the hunting season. Also included are the overall model concordance statistic with standard error (Model concordance (SE)) and results of the Wald test (model p-value) evaluating whether a model differs significantly from the null. MD = minimum distance, Pre 1 = previous 1 day average, Pre 3 = previous 3 day average, Pre 7 = previous 7 day average.

| Model | Covariate | Beta Coef. | Robust SE | p-value | HR (95% CI) | Model Concordance (SE) | Model p-value |
|---|---|---|---|---|---|---|---|
| Capture Mass + No. of Feeders in 95% Range | No. Feeders: 0-3 days | -0.07 | 0.04 | 0.05 | 0.93 (0.87-1.00) | 0.77 (SE < 0.01) | < 0.001 |
| | No. Feeders: 3-8 days | 0.22 | 0.06 | < 0.001 | 1.25 (1.12-1.40) | | |

| | | | | | | | |
|---|---|---|---|---|---|---|---|
| | No. Feeders: 8-36 days | 0.91 | 0.08 | < 0.001 | 2.49 (2.13-2.90) | | |
| Capture Mass + Pre 1 No. of Unique Feeders Visited Daily | Pre 1 No. Feeders Visited | 0.62 | 0.22 | 0.01 | 1.85 (1.20, 2.85) | 0.75 (SE = 0.04) | < 0.001 |
| Capture Mass + Pre 3 No. of Unique Feeders Visited Daily | Pre 3 No. Feeders Visited | 0.42 | 0.31 | 0.18 | 1.52 (0.82-2.81) | 0.71 (SE = 0.06) | < 0.001 |
| Capture Mass + Pre 7 No. Unique Feeders Visited Daily | Pre 7 No. Feeders Visited | 0.34 | 0.4 | 0.39 | 1.40 (0.71-3.05) | 0.70 (SE = 0.05) | < 0.001 |
| Capture Mass + Most Recent Roost MD | Roost MD | -0.8 | 0.1 | < 0.001 | 0.45 (0.37-0.55) | 0.78 (SE = 0.02) | < 0.001 |
| Capture Mass + Pre 1 Roost MD + Pre 1 Morning MD | Pre 1 Morning MD | -0.45 | 0.02 | < 0.001 | 0.64 (0.62-0.66) | 0.73 (SE = 0.05) | < 0.001 |
| | Pre 1 Roost MD | -0.11 | 0.37 | 0.77 | 0.90 (0.43-1.86) | | |
| | Pre 3 Morning MD | -0.87 | 0.01 | < 0.001 | 0.42 (0.41-0.43) | 0.75 (SE = 0.05) | < 0.001 |

| | | | | | | | |
|---|---|---|---|---|---|---|---|
| Capture Mass + Pre 3 Roost MD + Pre 3 Morning MD | Pre 3 Roost MD | -0.02 | 0.44 | 0.96 | 0.98 (0.41-2.32) | | |
| Capture Mass + Pre 7 Roost MD + Pre 7 Morning MD | Pre 7 Morning MD | -1.22 | 0.1 | < 0.001 | 0.30 (0.24-0.36) | 0.70 (SE = 0.03) | < 0.001 |
| | Pre 7 Roost MD | 0.45 | 0.41 | 0.27 | 1.56 (0.70-3.47) | | |
